# Derivation and characterization of an embryonic‑derived muscle progenitor cell line from Atlantic salmon (*Salmo salar*)

**DOI:** 10.64898/2026.04.13.718151

**Authors:** Kyle Naylor, Sophia Webb, Deepika Rajesh, Patrick J. Mee

## Abstract

Unlike mammals, teleost fish exhibit lifelong skeletal muscle growth, characterized by continued fiber hypertrophy and the formation of new muscle fibers maintained by a persistent progenitor cell population. However, the limited availability of stable muscle progenitor cell lines from commercially important species such as Atlantic salmon (*Salmo salar*) constrains mechanistic studies and emerging applications in cellular aquaculture. Here, we report the establishment and characterization of a novel embryonic-derived salmon muscle progenitor cell line, termed SsEC. These cells were derived from late embryonic stages and exhibited a spindle-shaped morphology, robust proliferative capacity, and sustained expansion beyond 30 passages under defined culture conditions. SsECs demonstrated a distinct extracellular matrix preference, with vitronectin supporting long-term maintenance and expansion. Molecular characterization confirmed stable expression of canonical myogenic markers, including *myf5* and *myod1*, while transcriptomic profiling revealed enrichment of genes associated with muscle development and sarcomere organization relative to a non-myogenic salmon cell line. Directed differentiation to muscle, using a two-step protocol, induced efficient formation of multinucleated myotubes expressing myosin heavy chain and sarcomeric α-actinin, with upregulation of key differentiation markers such as *myog* and *Tnnt3a*. Together, these findings establish SsECs as a robust *in vitro* model cell line for studying salmon muscle development and provide a novel platform for applications in aquaculture research and cellular seafood production.

## Introduction

Skeletal muscle constitutes most of the body mass in teleost fish and plays central roles in locomotion, metabolism, and aquaculture productivity. Unlike mammals, teleosts exhibit indeterminate growth, in which muscle mass increases throughout life via a combination of hypertrophy and sustained hyperplasia driven by resident myogenic progenitor cells (Gabillard et al. 2010). This growth strategy makes fish muscle an attractive system for studying vertebrate myogenesis as well as for the development of cell-based seafood platforms (Johnston et al. 2011; Joyce 2023; Rubio et al. 2019).

The molecular regulation of teleost myogenesis is governed by a conserved network of transcription factors, including the myogenic regulatory factors (MRFs) *myf5* (myogenic factor 5), *myod1* (myogenic differentiation 1), and *myog* (Myogenin), alongside paired-box proteins such as *pax*7 and *pitx2*, which together coordinate progenitor specification, proliferation, and differentiation (Rossi and Messina 2014; Ramírez de Acuña et al. 2022). These intrinsic programs are further modulated by environmental variables, including temperature, nutrition, and oxygen availability, which exert significant effects on muscle growth trajectories and phenotypic plasticity in aquaculture species (Macqueen et al. 2008; Johnston et al. 2011). Despite substantial progress, mechanistic understanding of teleost muscle development remains incomplete, particularly with respect to early progenitor populations and their behavior under in vitro conditions (Rossi and Messina 2014; Joyce 2023).

To address these gaps, vitro models have increasingly been employed as tools to dissect the cellular and molecular basis of myogenesis. However, compared with mammalian systems, fish muscle cell culture remains relatively underdeveloped and fragmented across species (Goswami et al. 2022; Iqbal et al. 2025). Early studies relied primarily on primary satellite cell cultures, which are limited by donor variability, finite proliferative capacity, and inconsistent differentiation outcomes (Gabillard et al. 2010). More recently, a small number of studies have reported the establishment of continuous or long-term propagating muscle cell lines from teleost species. These include myoblast or muscle-derived lines from Atlantic mackerel, black sea bream, grouper, seabass, and goldfish, demonstrating varying capacities for sustained proliferation and myogenic differentiation (Saad et al. 2023; Han and Gong 2024; Kong et al. 2021; Li et al. 2021; Xue et al. 2024). Notably, some continuous teleost muscle cell lines have also exhibited adipogenic-like plasticity under defined conditions, highlighting both their developmental flexibility and current limitations in lineage fidelity and maturation (Saad et al. 2023; Trace et al. 2024).

The development of stable fish muscle cell lines has gained renewed importance with the emergence of cellular aquaculture, an approach aimed at producing seafood through in vitro cell culture. Recent studies emphasize the need for well-characterized, scalable cell sources capable of sustained self-renewal and controlled differentiation to support the generation of structured muscle–fat tissue (Trace et al. 2024; Goswami et al. 2024). In this context, a continuous Atlantic salmon (*Salmo salar*) muscle cell line with both myogenic differentiation and inducible adipogenic transdifferentiation capacity has recently been reported, representing a major advance for a commercially important species while also underscoring persistent challenges in achieving robust functional maturation and long-term lineage stability (Wei et al. 2025).

In addition to lineage stability, the developmental origin of cell lines is increasingly recognized as a key determinant of proliferative capacity, plasticity, and long-term utility (Musialak et al. 2024). Most existing fish muscle models are derived from juvenile or adult tissues, whereas embryonic-derived progenitor populations may offer enhanced developmental plasticity and suitability for extended culture (Rossi and Messina 2014). However, robust embryonic-derived muscle progenitor cell lines from commercially important teleosts such as Atlantic salmon remain scarce, limiting both fundamental studies of early myogenesis and the development of scalable platforms for aquaculture biotechnology and cultivated seafood applications.

In this study, we describe the generation and characterization of an embryonic Atlantic salmon muscle progenitor cell line (SsEC) and compare its properties with those of a well-established Atlantic salmon kidney cell line (ASK). Unlike ASKs, SsECs display a distinct extracellular matrix preference during early embryonic stages, sustain long-term proliferation, and retain reproducible differentiation into sarcomere-bearing myotube lineages in vitro. These features establish SsECs as a valuable model for investigating early teleost myogenesis and embryonic muscle lineage specification.

## Materials and Methods

### Fish Source and Embryo Collection

Atlantic salmon (*Salmo salar*) fertilized ova (StofnFiskur Salmoselect, diploid mixed-sex) were obtained from Benchmark Genetics. Embryos were incubated at ∼4.5°C in aerated freshwater until the desired developmental stage. Eyed-stage embryos were collected at ∼370 Biological Degree Days of Growth (B-DDG) (B-DDG; calculated as incubation temperature × days), post-fertilization depending on incubation temperature, and blastula-stage embryos at ∼40-48 B-DDG post-fertilization; staging was confirmed morphologically (optic cup for eyed-stage; hollow-sphere morphology for blastula) (Musialak et al. 2024)

### Isolation and culture of Salmon Embryo Derived Cells

Embryos were rinsed once in 70% ice-cold ethanol and three times in Dulbecco’s Phosphate Buffered Saline, no calcium, no magnesium (Gibco, 14190144) (DPBS)containing 100 U/mL Penicillin-Streptomycin (Gibco, 15070063) and 2.5 µg/mL Amphotericin B (Gibco, 15290018). Embryos were homogenized in sterile DPBS and the homogenate was filtered sequentially through 200 µm and 100 µm meshes to remove chorion and debris, then centrifuged at 300 × g for 4 min at 4°C. Finally, the pellet was resuspended in complete Salmon Embryo Derivation (ED) medium (DMEM/F-12 [Thermo Fisher, 15070-063], 10% FBS [VWR, 97068-085], 100 U/mL penicillin-streptomycin [Thermo Fisher, 15070-063], 1 × Non-Essential Amino Acids [NEAA] [Thermo Fisher, 11140050], 1 mM Sodium pyruvate [Thermo Fisher, 11360070], 27.5 µM 2-mercaptoethanol [Thermo Fisher, 31350010], 15 mM HEPES [Merck, H0887], 20 µg/mL FGF-2 [PeproTech, 100-18B]).

Following resuspension, cells were seeded into T-25 tissue culture flasks, pre-coated with either 0.2% Gelatin (Sigma-Aldrich, G1393), 0.5 µg/cm^2^ Laminin-511 (Sigma-Aldrich, CC160), or 0.5 µg/cm^2^ Vitronectin (Thermo Fisher, A14700). Cells were maintained at 20°C with 5% CO_2_ with media being replaced every 48 h. Initial attachment occurred within 2–7 days and cultures reached confluency by ∼day 14. Cells were passaged with Accutase (Merck, A6964), maintaining a seeding density of 3 × 10^3^/cm^2^ at each passage. Cells were successfully expanded for >30 passages without loss of progenitor phenotype.

All cell banks were tested for microbial contamination by inoculation into Thioglycollate Medium (VWR, 621051ZA) and Tryptone Soy Broth (Millipore, STBMTSB12) and monitored for growth over a 14-day incubation period. Mycoplasma contamination was assessed using the Venor GeM qEP Mycoplasma Detection Kit (MinervaBiolabs, 11-9250) following the manufacturer’s protocol. Species identity was confirmed by sequencing of PCR products, RNA alignment identity to model genome, and karyotyping (VHL genetics).

### Culture of Atlantic Salmon Kidney (ASK) Cells

ASK cells (ATCC CRL-2747 or equivalent) were maintained in Leibovitz’s L-15 medium (Sigma-Aldrich, L4386) supplemented with 10% FBS, 2 mM L-glutamine (Gibco, A29168-01), and 100 U/mL penicillin–streptomycin at 20°C in ambient air (no CO_2_), consistent with provider guidelines (ATCC; Cellosaurus). Cells were seeded at 1 × 10^4^ cells/cm^2^, in vessels precoated with 0.2% gelatin, and subcultured every 7–10 days using a cell scraper; media were refreshed every 2-3 days (ATCC, 2026; Cellosaurus, 2025).

### Myogenic Differentiation of SsECs

Directed differentiation was performed using a modified two-step protocol previously reported by Sanaki-Matsumiya et al. 2024.

In the first step, presomitic mesoderm (PSM) induction (days 0–2), cells were cultured in either NBFR medium (Condition 1) or Salmon ED medium (Condition 2) supplemented with the GSK-3 inhibitor CHIR99021 dihydrochloride (10 μM; Axon Medchem, 2435) to activate canonical WNT signaling, the ALK4/5/7 inhibitor SB431542 (10 μM; Selleck Biotechnology Ltd, S1067) to suppress TGF-β signaling, the BMP inhibitor DMH1 (2 μM; Sigma/Merck, D8496), and salmon bFGF (20 ng/mL; Qkine Qk102-FG). This combination was intended to promote mesodermal commitment while preventing premature differentiation.

In the second step, myogenic differentiation (days 3–17), two growth factor–supplemented media sequences were evaluated. In Condition 1 (C1), PSM-primed cells were transferred to Muscle Medium 1 (MM1: DMEM/F-12 supplemented with 15% Knockout Serum Replacement (KSR) [Gibco, 10828028], 10 ng/mL IGF-1 [PeproTech, 100-11], 10 ng/mL HGF [PeproTech, 100-39], and 20 ng/mL salmon bFGF) for 4 days, followed by Muscle Medium 2 (MM2: DMEM/F-12 supplemented with 15% KSR, 10 ng/mL IGF-1, and 10 ng/mL HGF, without bFGF) until day 17. In Condition 2 (C2), PSM-primed cells were transferred to Muscle Salmon Medium 1 (MS1: Salmon ED medium supplemented with 5% FBS, 10 ng/mL IGF-1, 10 ng/mL HGF and 20 ng/mL salmon bFGF) for 4 days, followed by Muscle Salmon Medium 2 (MS2: Salmon ED medium supplemented with 5% FBS, 10 ng/mL IGF-1, and 10 ng/mL HGF, without bFGF) until day 17. Media was changed every other day in both conditions.

### RNA Extraction and Quantitative PCR (qPCR)

Total RNA was extracted using the RNeasy Mini Kit (Qiagen, 74104) according to the manufacturer’s instructions. RNA integrity was assessed by NanoDrop and Bioanalyzer where indicated. cDNA was synthesized with the AffinityScript™ Multiple Temperature Reverse Transcriptase (Agilent Technologies, 600105) according to the manufacturer’s instructions. and qPCR performed on a QuantStudio 3 (Applied Biosystems) using SYBR Green. Primers targeted *myf5* (myogenic factor 5), *tpm1* (tropomyosin 1), *acta1* (actin alpha 1), *myod1* (myogenic differentiation 1), *myog* (myogenin), *tnnt3a* (troponin T type 3a), *vwf* (von Willebrand factor), *Pecam1* (platelet and endothelial cell adhesion molecule 1), and *kdr* (kinase insert domain receptor), with *ube2l3* (ubiquitin-conjugating enzyme E2 L3) as the endogenous reference gene (primer sequences are provided in Table S1). Relative expression was calculated by the ΔΔCt method. All reactions were performed in biological triplicate.

### Bulk RNA-Sequencing Analysis

Total RNA was isolated from SsECs at passages 8 and 32 and from ASK cells, with three independent biological replicates per condition. RNA integrity was assessed using an Agilent Bioanalyzer, and only samples with an RNA integrity number (RIN) ≥ 8 were used for library preparation. Sequencing libraries were generated using the TruSeq Stranded mRNA Library Preparation Kit (Illumina) according to the manufacturer’s instructions. Libraries were sequenced on an Illumina NovaSeq 6000 platform to produce 150 bp paired-end reads, with an average depth of approximately 30 million reads per sample. Library preparation and sequencing were performed by GENEWIZ (Azenta Life Sciences).

Raw sequencing reads were assessed for quality using FastQC (v0.12.1). Adapter sequences and low-quality bases were trimmed using Cutadapt (v3.2), removing reads with a Phred quality score <30, reads shorter than 50 bp after trimming, and reads containing ambiguous nucleotides. High-quality reads were aligned to the Atlantic salmon reference genome (Ensembl Ssal_v3.1; GCA_905237065.2) using STAR aligner (v2.7.11a) with default parameters. Gene-level read counts were generated using featureCounts (v2.0.6) (Liao et al. 2014) based on Ensembl gene annotations.

Differential gene expression analysis was performed in R using edgeR (v4.2.2). Lowly expressed genes were filtered prior to analysis, and normalization was conducted using the trimmed mean of M-values (TMM) method. Dispersion estimates were calculated using a generalized linear model framework, and differential expression between groups was assessed using quasi-likelihood F-tests. Genes with a false discovery rate (FDR) < 0.05 and an absolute log_2_ fold change ≥ 1.5 were considered significantly differentially expressed.

Raw and processed RNA-seq data are available from Roslin Technologies Ltd servers. Access to these datasets will be provided upon request to the corresponding author.

### Pathway Enrichment Analysis

Enrichment of differentially expressed genes (DEGs) was evaluated using Gene Ontology (GO) and Kyoto Encyclopedia of Genes and Genomes (KEGG) over-representation analyses implemented in the R package clusterProfiler (v4.12.6; Wu et al. 2021). GO terms and KEGG pathway analyses using the hypergeometric test were performed to identify significantly enriched pathways. Pathways with false discovery rates (FDR) *P* < 0.05 were deemed significant.

### Immunocytochemistry (ICC)

Cells were fixed in 4% paraformaldehyde (Thermo Fisher Scientific, J61889.AK) (15 min, room temperature), cells were washed with PBST (0.2% Tween20 (Fisher BioReagents, BP337-100) in PBS) for 5 min and blocked with 3% normal Bovine serum albumin (Sigma Aldrich, A1470) in PBST (1 h, room temperature). Cells were then incubated overnight at 4°C with primary antibodies against sarcomeric α-actinin (SAA; clone EA-53, Abcam, cat. no. ab9465, 1:1000) and myosin heavy chain (MHC; clone MF20, Bio-techne, 1:300). After three PBS washes, cells were incubated with Alexa Fluor 488-conjugated secondary antibody (Thermo Fisher Scientific, A-21202, 1:1000) for 1 h at room temperature in the dark. Nuclei were counterstained with NucBlue (Invitrogen, R37605) (1 μg/mL, 15 min). Images were acquired on a Nikon Eclipse TS2 inverted fluorescence microscope.

### Statistical Analysis

Unless stated otherwise, experiments were performed in biological triplicate. Data are presented as mean ± SD. Statistical significance was assessed by one-way ANOVA with Tukey’s *post hoc* test (GraphPad Prism v9), with *P* < 0.05 considered significant. Specific *P* values are reported in figures. GraphPad Prism (v9) and R Studio (v4.4.0) were used for data analysis and visualization.

## Results

### Extracellular matrix composition determines successful derivation and long-term expansion of salmon embryonic progenitors

To establish defined culture conditions capable of supporting the derivation and sustained expansion of embryonic salmon muscle progenitors, we first evaluated the influence of extracellular matrix (ECM) composition on cell attachment and growth. Cells dissociated from late eyed-stage Atlantic salmon embryos exhibited pronounced substrate-dependent attachment and proliferation behaviors. When plated on gelatin, cells attached transiently but showed minimal proliferation, with progressive detachment and culture failure by approximately 30 days (Figure 2A). Cultures established on laminin-coated surfaces displayed improved initial attachment and proliferation; however, growth was not sustained beyond ∼30 days, and cells consistently failed to reattach following enzymatic passaging (Figure 2B).

In contrast, vitronectin supported robust initial cell attachment, sustained proliferation, and serial passaging, enabling continuous expansion of embryonic-derived salmon progenitors for more than 60 days in culture and beyond 30 passages (Figures 1–3). Cells maintained stable morphology and growth characteristics across passages under vitronectin-supported conditions. Collectively, these findings identify vitronectin as a critical ECM component for the successful derivation, long-term maintenance, and expansion of embryonic salmon muscle progenitor cells (SsECs).

**Figure 1.**
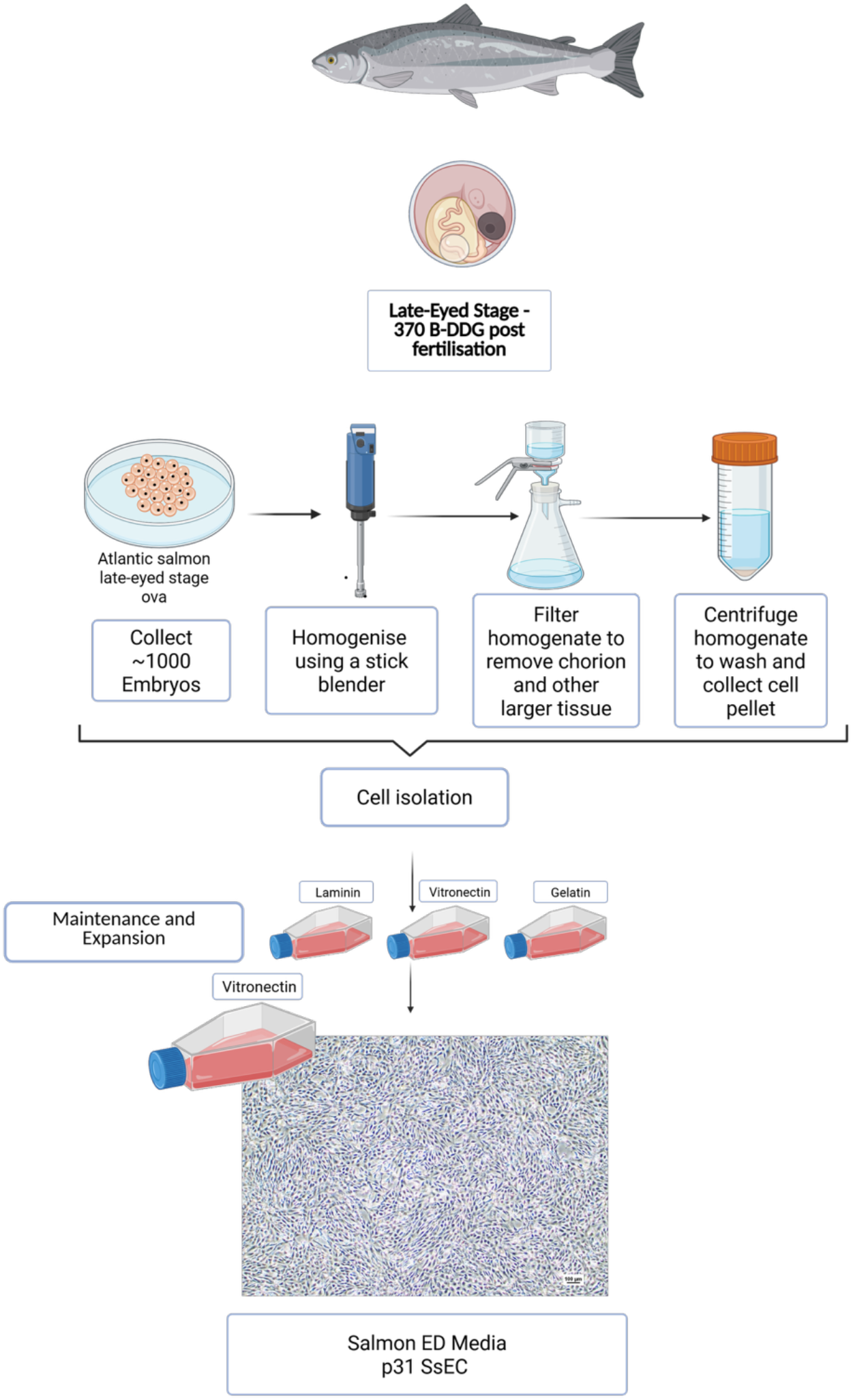
Derivation of SsECs from late eyed-stage Atlantic salmon (*Salmo salar*) embryos. Schematic overview of embryonic tissue dissociation and seeding of the resulting cell suspension onto candidate extracellular matrix substrates (laminin, vitronectin, gelatin) in Salmon ED medium. Vitronectin supported sustained attachment and long-term expansion of SsECs. Representative phase-contrast image shows SsECs cultured on vitronectin at passage 31.

**Figure 2.**
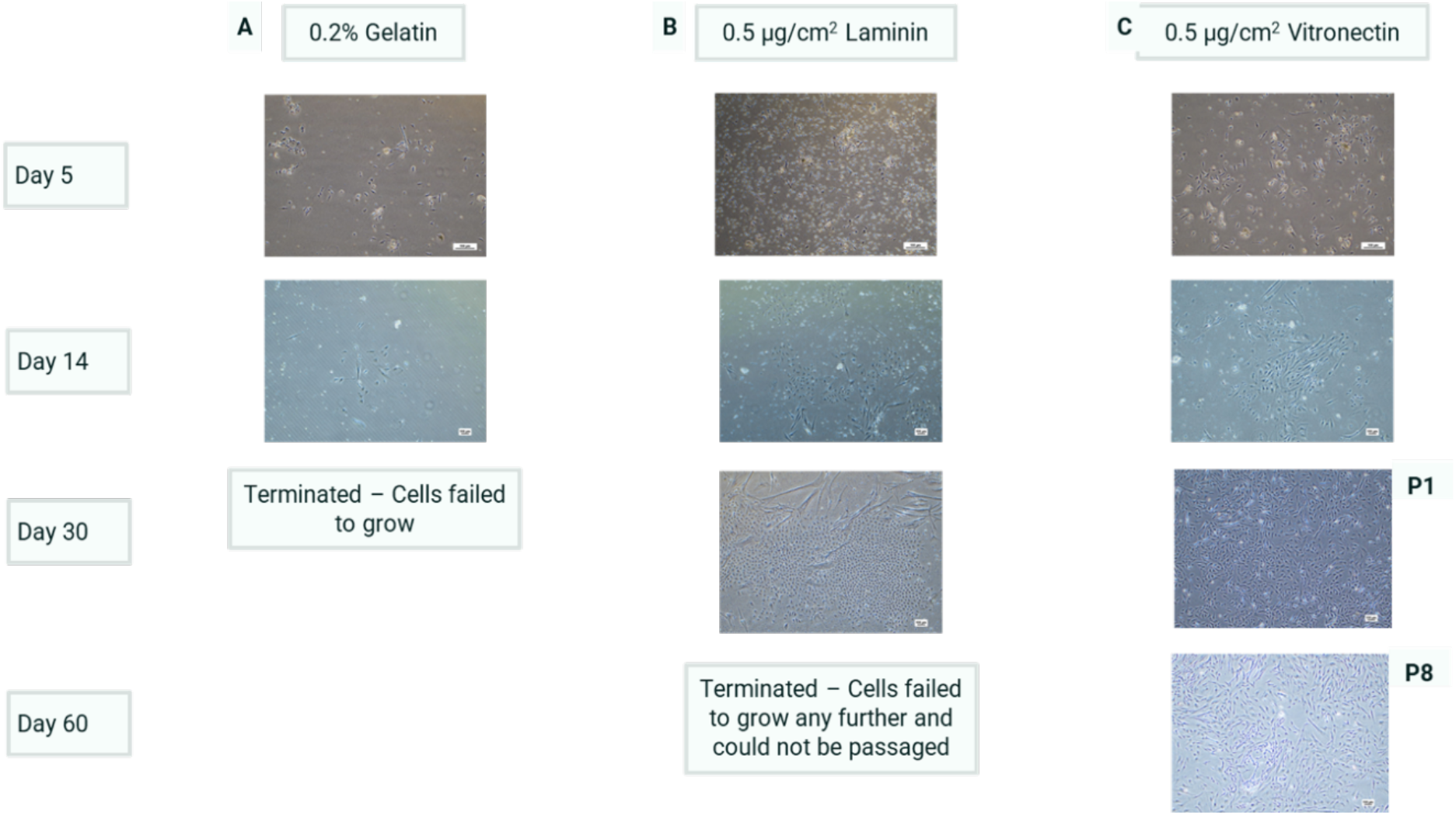
Substrate dependence of SsEC derivation and early expansion from late eyed-stage Atlantic salmon (*Salmo salar*) embryos. Cells were cultured on gelatin, laminin, or vitronectin under identical conditions. (A) Gelatin supported transient attachment, followed by loss of proliferation and culture failure by ∼30 days. (B) Laminin permitted initial attachment and proliferation, but cultures failed to sustain growth or reattach after passaging. (C) Vitronectin supported robust attachment, continued proliferation, and serial passaging beyond 60 days.

### SsECs exhibit accelerated growth kinetics and distinct morphology compared to ASK cells

To contextualize the growth behavior of SsECs, these cells were compared to an existing and widely utilized Atlantic salmon kidney (ASK) cell line to compare proliferation and morphology. Both cell types were cultured under their respective standard conditions, and growth kinetics were assessed by measuring viable cell numbers over time to estimate population doubling times, while morphology was evaluated by phase-contrast microscopy.

SsEC cultures displayed a markedly different morphology compared to ASK cells. SsECs exhibited an elongated, spindle-shaped morphology consistent with a progenitor-like phenotype, whereas ASK cells showed a flatter, more epithelial-like appearance typical of kidney-derived cell lines (see Figure 3B–C).

**Figure 3.**
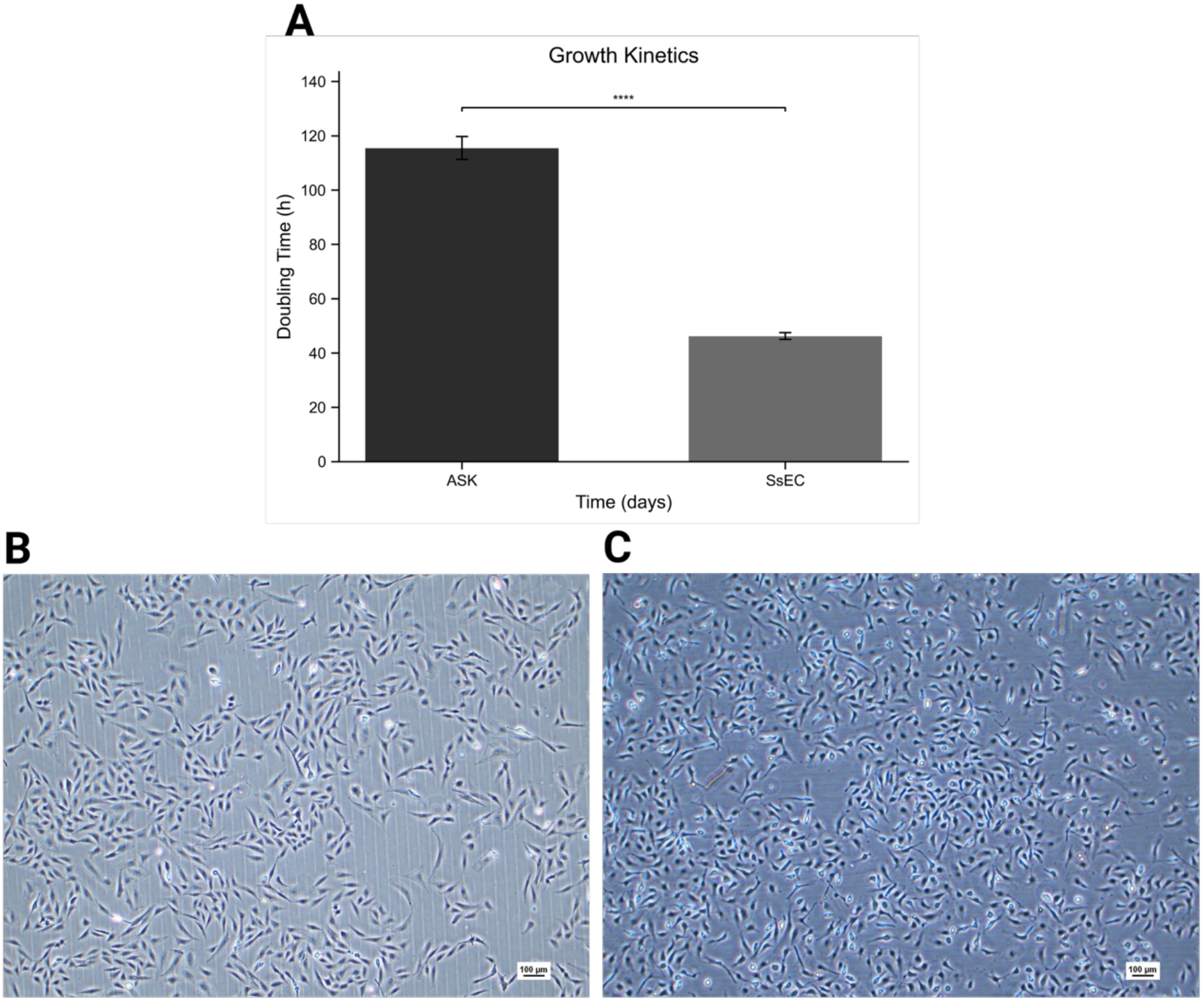
Growth kinetics and morphology of SsECs and ASK cells. (A) Population doubling times of ASK cells and SsECs under standard culture conditions. SsECs exhibited a significantly shorter doubling time than ASK cells (*P* < 0.0001, unpaired *t*-test; mean ± SD, *n* = 3). (B, C) Representative phase-contrast images showing spindle-shaped, progenitor-like morphology in SsECs (B) and flattened epithelial/endothelial-like morphology in ASK cells (C).

Quantitative analysis of growth kinetics revealed that SsECs proliferated substantially faster than ASK cells, with an estimated population doubling time of approximately 54 hours for SsECs compared to ∼120 hours for ASK cells (Figure 3A). This difference indicates an enhanced proliferative capacity in SsECs under the conditions tested.

Together, these results demonstrate that SsECs are both morphologically and kinetically distinct from a well-established non-myogenic salmon cell line, supporting their classification as a rapidly expanding progenitor population and reinforcing their suitability for downstream molecular and functional analyses.

### Transcriptomic profiling identified a musculoskeletal program in SsEC

To determine whether the phenotypic and proliferative differences observed in SsECs relative to ASK cells were reflected at the molecular level, we performed bulk RNA-sequencing to compare the global transcriptional profiles of the two cell populations (passage 8) and late (passage 32) culture stages and compared with Atlantic salmon kidney (ASK) cells as a non-myogenic reference line.

At the global transcriptome level, principal component analysis (PCA) revealed a clear separation between SsEC and ASK samples, with the first principal component (PC1) accounting for 84.27% of the total variance (Figure 4A). The second and third components explained substantially less variance (4.10% and 3.98%, respectively), indicating that the dominant source of transcriptional variation was cell-type identity rather than passage number or technical variability. Notably, SsEC samples clustered closely across passages, supporting the stability of the SsEC transcriptional program during long-term culture.

**Figure 4.**
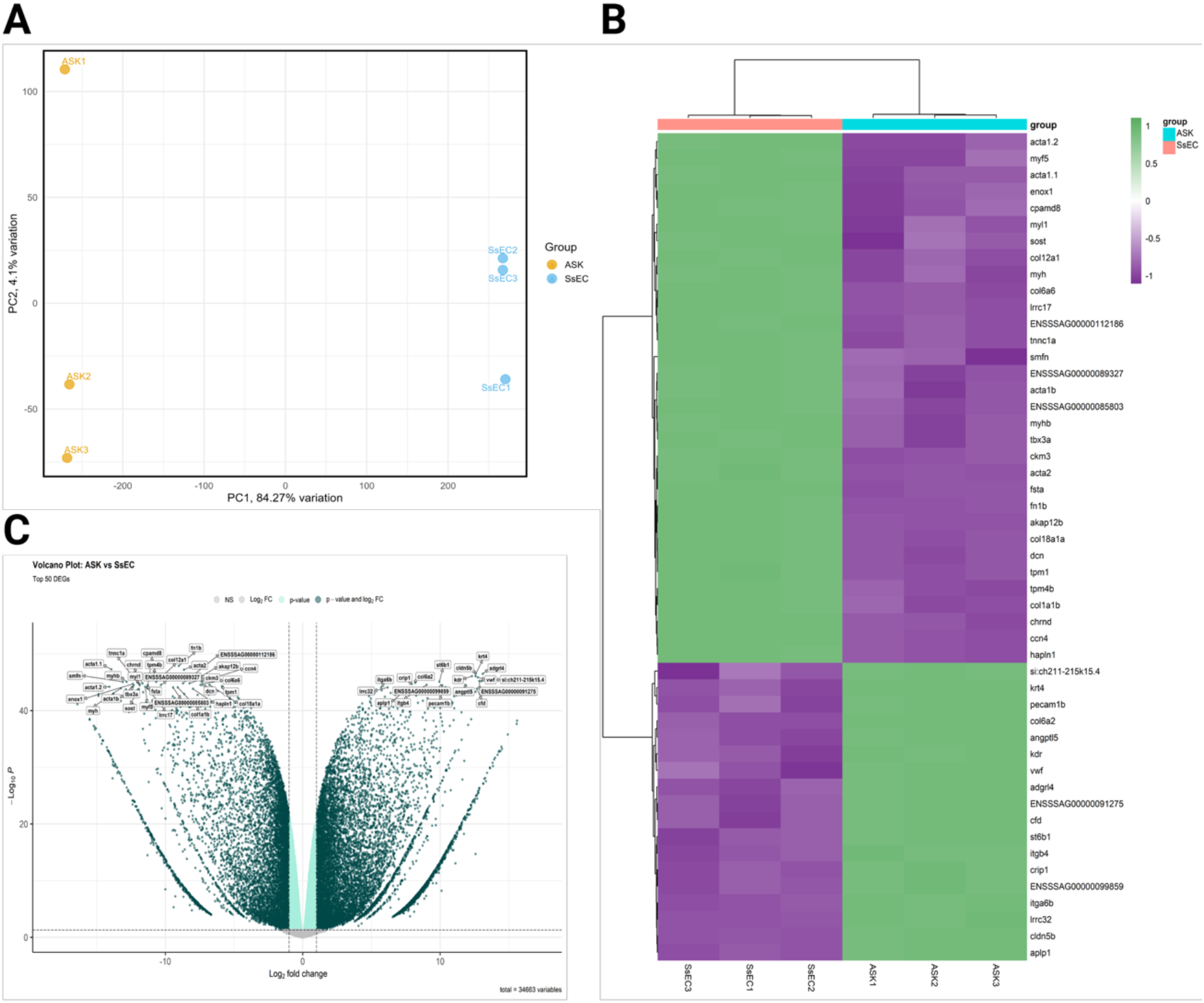
Transcriptomic profiling of SsECs and ASK cells. (A) Principal component analysis of RNA-sequencing data showing clear separation between SsEC and ASK samples, with PC1 and PC2 accounting for 84.27% and 4.10% of total variance, respectively. (B) Volcano plot of differentially expressed genes between SsECs and ASK cells (adjusted *P* < 0.05; absolute log_2_ fold change ≥ 1.5). (C) Heatmap of the top 50 differentially expressed genes with hierarchical clustering, illustrating enrichment of myogenic genes in SsECs (e.g. *myf5, acta1, tpm1, myhb*) and endothelial-associated genes in ASK cells (e.g. *vwf, kdr, pecam1b*). RNA-sequencing was performed using three replicates per cell type.

To identify the molecular features underlying this separation, differential expression analysis (DEG) was performed between SsECs and ASK cells using edgeR following read alignment with STAR and summarization with featureCounts. Applying a false discovery rate (FDR) threshold of < 0.05 and an absolute log_2_ fold change (|log_2_FC|) ≥ 1.5, a large set of genes was identified as being differentially expressed between SsECs and ASK cells. Of which 14,782 genes were upregulated in ASK, 13,749 genes were up regulated in SsEC and Volcano plot visualization highlighted pronounced enrichment of genes associated with myogenesis, muscle structure, and cytoskeletal organization in SsECs, whereas genes upregulated in ASK cells were predominantly associated with endothelial identity and kidney-derived lineages (Figure 4C). A heatmap of the top 50 differentially expressed genes further emphasized this transcriptional dichotomy, with coordinated upregulation of muscle-associated genes in SsECs and endothelial markers in ASK cells (Figure 4B).

Functional enrichment analysis of genes upregulated in SsECs reinforced the transcriptomic distinction between the two cell populations. Gene Ontology (GO) over-representation analysis identified significant enrichment of biological processes related to muscle development, sarcomere organization, myofibril assembly, and cytoskeletal organization. Consistently, Kyoto Encyclopaedia of Genes and Genomes (KEGG) pathway analysis highlighted pathways linked to muscle contraction and actin–myosin interactions. Together, these analyses indicate that SsECs are transcriptionally enriched for a musculoskeletal program that is absent in ASK cells.

To validate the RNA-sequencing findings and examine lineage marker expression across extended passaging, targeted RT–qPCR analyses were performed for a panel of myogenic and endothelial genes. SsECs exhibited significantly higher expression of structural and differentiation-associated muscle markers, including *acta1, tpm1*, and *tnnt3a*, relative to ASK cells at both passage 8 and passage 32 (Figures 5). Expression of the myogenic progenitor marker *myf5* was maintained and elevated at passage 32 compared with passage 8, consistent with preservation of progenitor features during long-term expansion. In contrast, expression of more differentiated muscle markers, including *myl1, tpm1*, and *acta1*, was higher at passage 8 than at passage 32, indicating limited spontaneous differentiation during extended culture (Figure 5).

**Figure 5.**
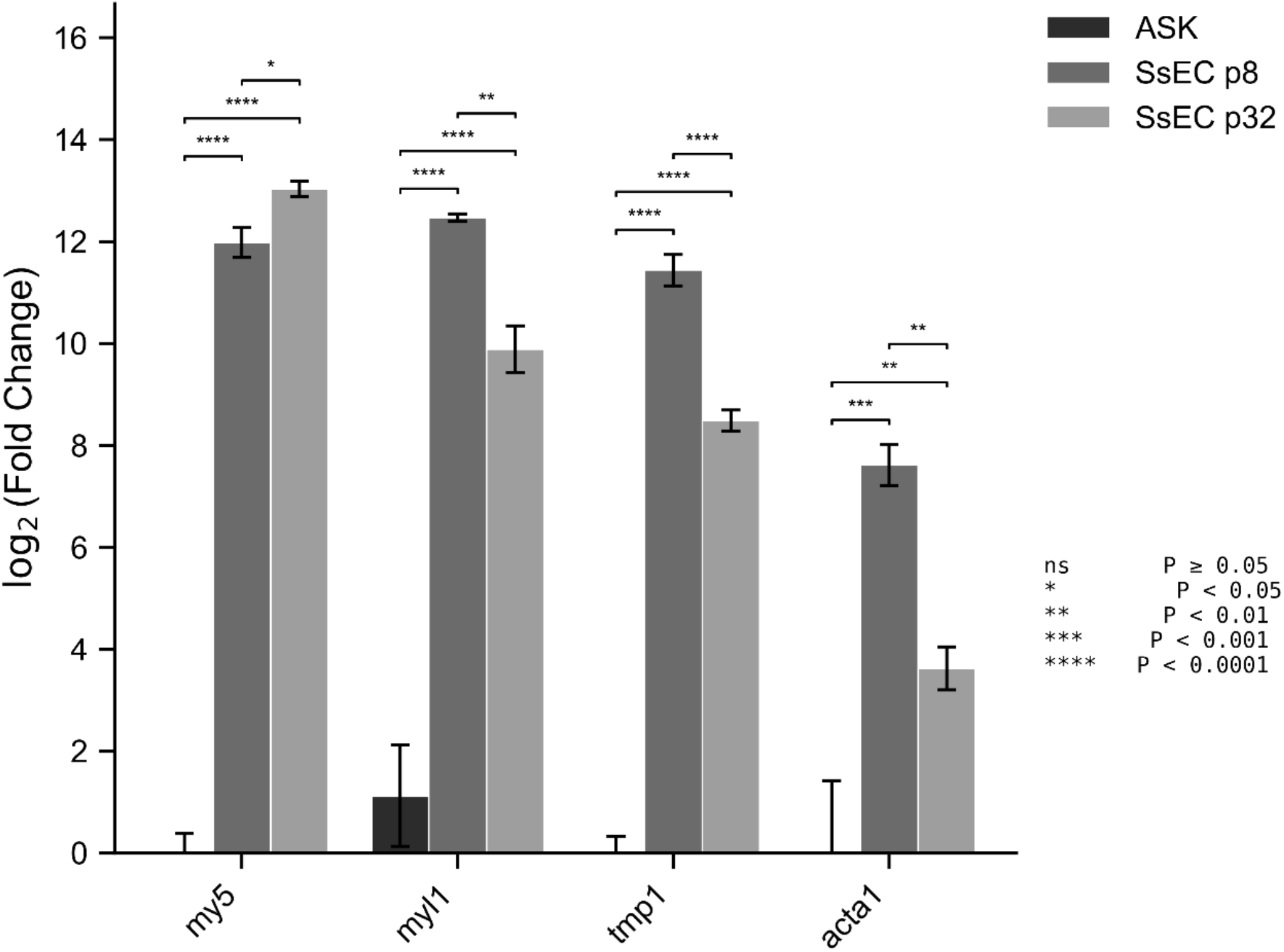
Quantitative PCR validation of musculoskeletal marker expression in SsECs and ASK cells. Relative log_2_(fold change) expression of *myf5, myL1, tpm1*, and *acta1* was assessed in SsECs and ASK cells at passage 8 (p8) and passage 32 (p32). All markers were expressed at higher levels in SsECs compared with ASK cells at both passages. Data are presented as mean ± SD (*n* = 3). Statistical significance was determined by two-way ANOVA with Tukey’s post hoc test.

Endothelial-associated genes, including *vwf, pecam1*, and *kdr*, were robustly expressed in ASK cells but were negligible or undetectable in SsECs at both passages examined (Figure 6). Although low-level *vwf* and *kdr* expression was detectable in late-passage SsECs, transcript abundance remained several orders of magnitude lower than that observed in ASK cells. These targeted expression analyses corroborated the RNA-sequencing results and confirmed that SsECs maintain a stable, lineage-restricted myogenic transcriptional profile across prolonged *in vitro* expansion, distinct from non-myogenic salmon cell lines.

**Figure 6.**
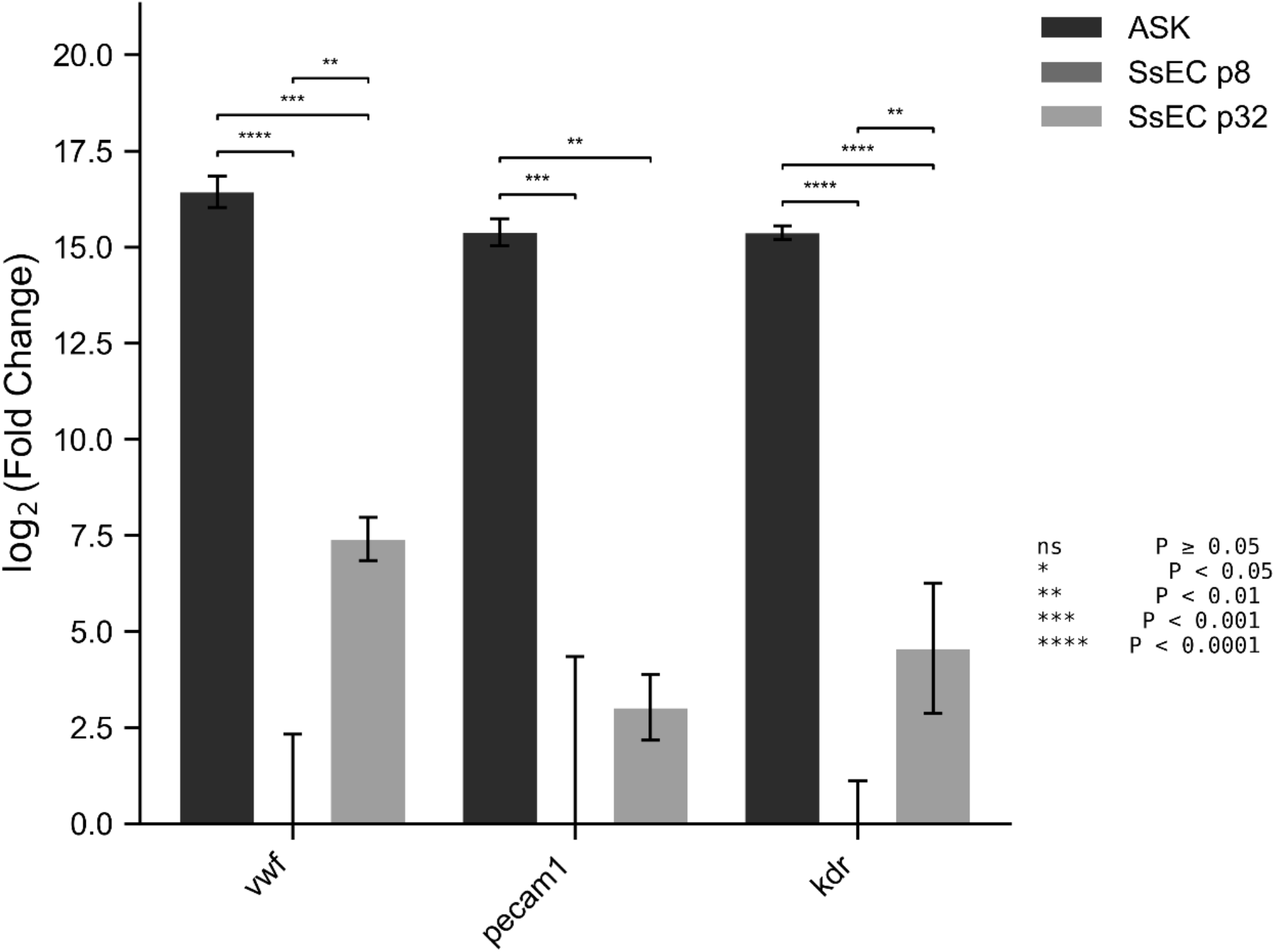
Quantitative PCR validation of endothelial marker expression in SsECs and ASK cells. Relative log_2_(fold change) expression of *vwf, pecam1* was assessed in SsECs and ASK cells at passage 8 (p8) and passage 32 (p32). Both markers were expressed at higher levels in ASK cells compared with SsECs at both passages. Data are presented as mean ± SD (*n* = 3). Statistical significance was determined by two-way ANOVA with Tukey’s post hoc test.

Collectively, these transcriptomic analyses demonstrate that SsECs possess a stable and distinct molecular identity that is strongly enriched for myogenic and musculoskeletal gene programs when compared with an established non-myogenic salmon cell line. These data provide a molecular framework supporting the classification of SsECs as embryonic-derived muscle progenitor cells and establish a rationale for targeted validation of lineage-specific markers by quantitative PCR in the subsequent section.

### Directed differentiation generated sarcomere-forming myotubes with robust MHC and α-actinin expression

To assess the myogenic differentiation capacity of SsECs and to compare fish-adapted serum containing media and serum-free differentiation environments, cells were subjected to a staged, two-step differentiation protocol designed to mimic key features of vertebrate myogenesis (Figure 7). In the first phase (days 0–2), SsECs were exposed to a presomitic mesoderm (PSM) induction regimen incorporating canonical WNT activation (CHIR99021) together with inhibition of TGF-β (SB431542) and BMP (DMH1) signaling. This signaling combination was applied under two basal media contexts namely, a serum-free NBFR formulation (Condition 1) or Salmon Embryo Derivation (ED) medium (Condition 2). This shared induction phase was intended to promote mesodermal commitment while preventing premature differentiation.

**Figure 7.**
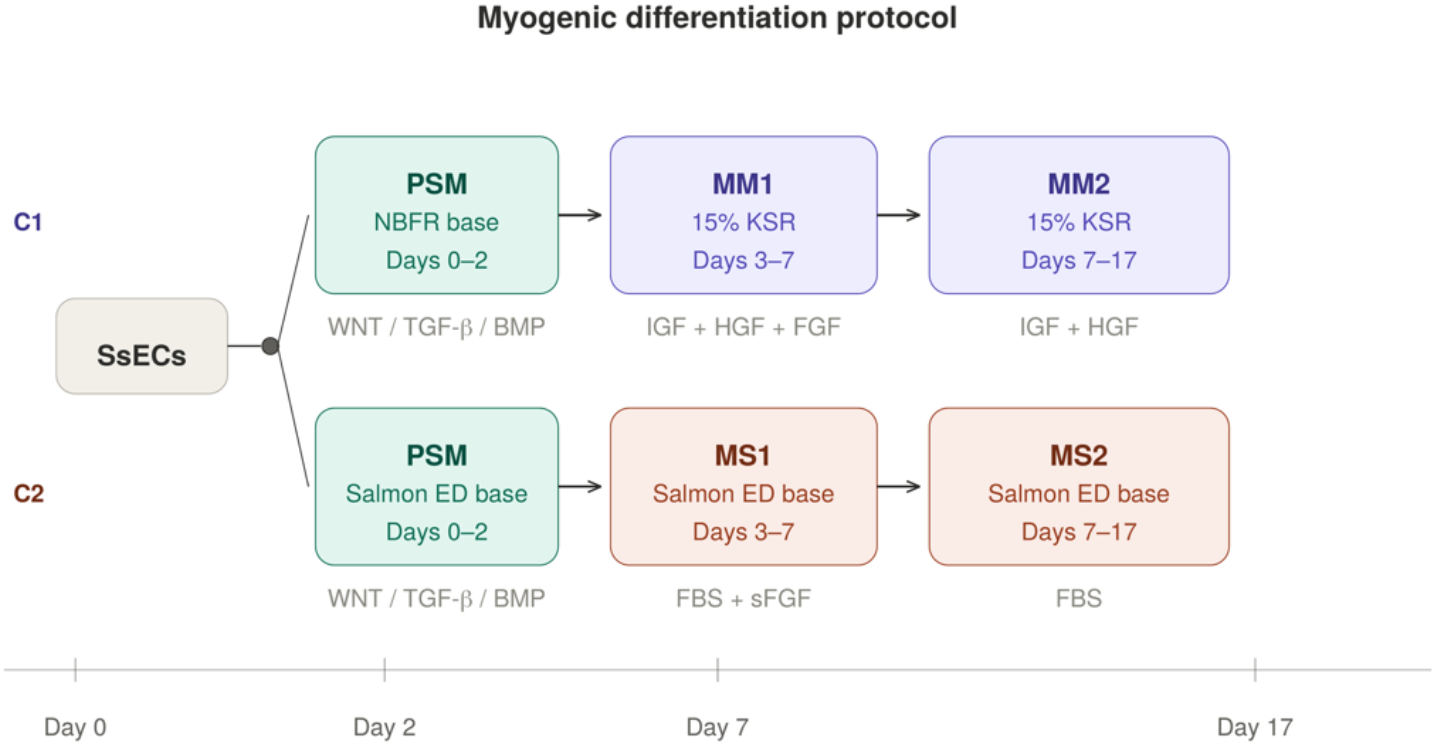
Schematic overview of the two-step myogenic differentiation protocol used for SsECs. Cells were first primed toward a presomitic mesoderm (PSM)-like state (days 0–2) through modulation of WNT, TGF-β, and BMP signaling, followed by transfer to growth factor– supplemented media to promote myogenic differentiation (days 3–17). Two differentiation conditions were compared: a serum-free formulation (Condition 1; MN1/MN2) and a low-serum Salmon ED-based formulation (Condition 2; MS1/MS2). bFGF was included transiently during early differentiation and withdrawn during terminal myogenic maturation.

Following PSM induction, cells were transitioned to differentiation media for progression toward terminal myogenesis (days 3–17), using distinct growth factor formulations designed to balance continued proliferation with differentiation (Figure 7). In Condition 1, PSM-primed cells were cultured in KSR-based muscle media (MN1 followed by MN2), whereas Condition 2 employed Salmon ED-based media supplemented with low-serum and salmon-specific growth factors (MS1 followed by MS2). In both conditions, bFGF was included transiently during the early differentiation phase and subsequently withdrawn to promote cell-cycle exit and myotube formation.

Quantitative RT–qPCR analysis at day 17 revealed that both conditions produced substantial upregulation of myogenic markers relative to the ASK control cell line (Figure 8). The myogenic commitment factor *myod1* was strongly induced under both formulations, with C1 (MN1/MN2) and C2 (MS1/MS2) reaching approximately 3,400-fold and 3,000-fold increases over ASK, respectively (ANOVA *P* < 0.0001; Figure 8A). Both conditions also significantly exceeded undifferentiated SsECs, which showed only modest *myod1* expression (∼400-fold over ASK), confirming that the differentiation protocol actively drives myogenic commitment beyond the basal progenitor state.

**Figure 8.**
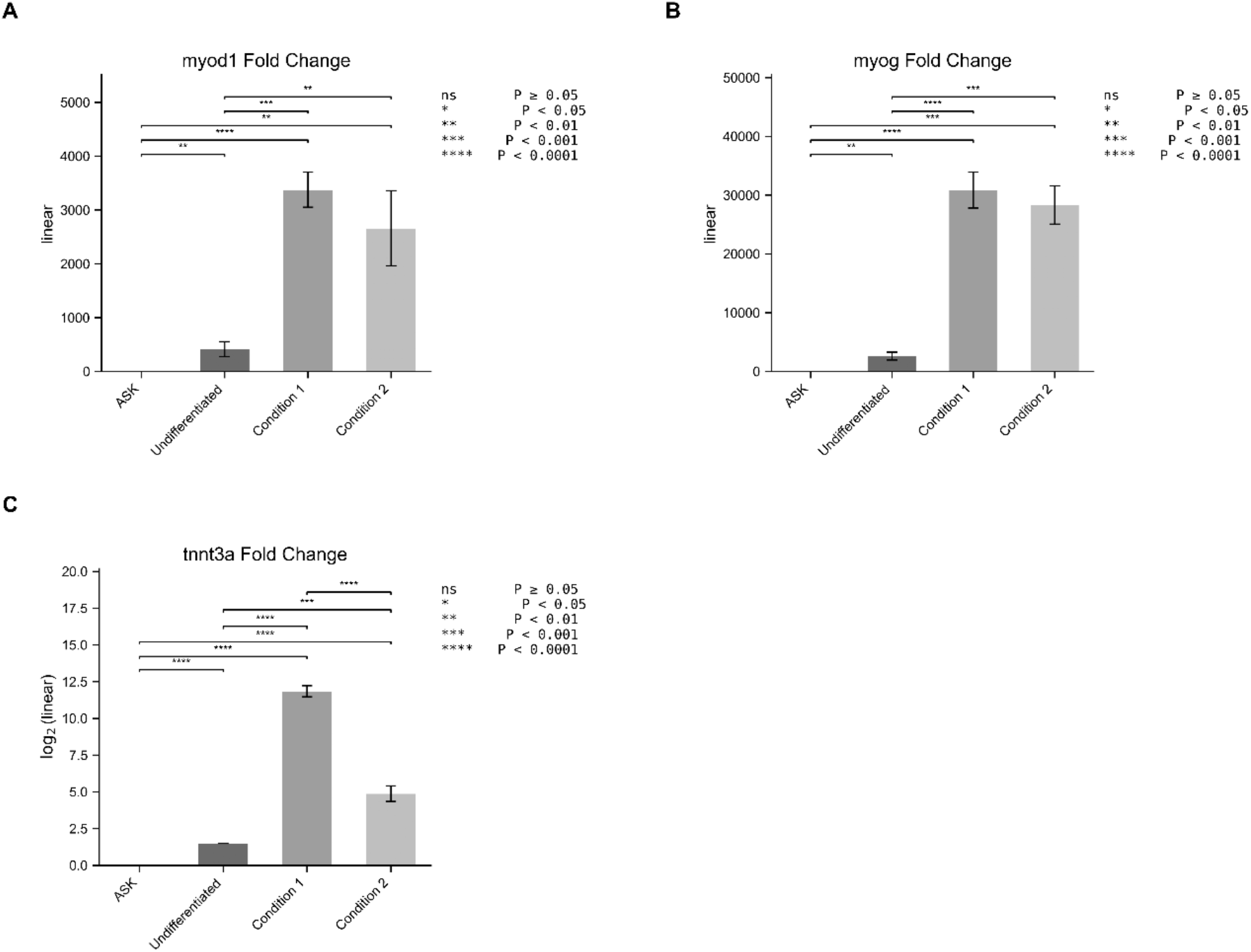
Quantitative PCR analysis of myogenic differentiation marker expression following directed induction. Relative expression of *myod1* (A), *myog* (B), and *Tnnt3a* (C) was assessed in ASK cells, undifferentiated SsECs, and SsECs differentiated under Condition 1 (C1; MN1/MN2) or Condition 2 (C2; MS1/MS2). Expression levels are shown as fold change relative to ASK for *myod1* and *myog*, and as log_2_(fold change) for *tnnt3a*. Data are presented as mean ± SD (*n* = 3). Statistical significance was determined by one-way ANOVA with Tukey’s post hoc test (*P* < 0.05).

Similarly, expression of *myog*, a transcription factor required for terminal differentiation and myotube fusion, was markedly elevated under both conditions, with C1 (MN1/MN2) and C2 (MS1/MS2) reaching approximately 30,000-fold and 28,000 increases over ASK respectively (ANOVA *P* = 0.0017; Figure 8B). This further suggests that both differentiation medias can effectively support the later stages of myogenic commitment in Atlantic salmon progenitors.

In contrast, the late sarcomeric marker *tnnt3a* showed C1 (MN1/MN2) produced substantially higher *tnnt3a* expression (∼11.8 log_2_ fold change over ASK) than C2 (MS1/MS2; ∼4.9 log_2_ fold change), with highly significant differences between all groups (ANOVA *P* < 0.0001; Figure 8C). As *tnnt3a* encodes a structural component of the sarcomeric thin filament, its preferential upregulation under the KSR-based formulation suggests that the serum-free environment may favor later stages of myofibrillogenesis, potentially by reducing mitogenic signaling that could otherwise delay exit from the cell cycle and sarcomere assembly.

Immunocytochemistry at day 15 showed strong staining for sarcomeric alpha-actinin (Figure 9) and myosin heavy chain (Figure 10), with visible alignment of contractile structures in the most effective condition. Lower-induction conditions yielded weaker staining and reduced cell density, suggesting that a balance between cell growth and differentiation was critical for efficient myotube formation.

**Figure 9.**
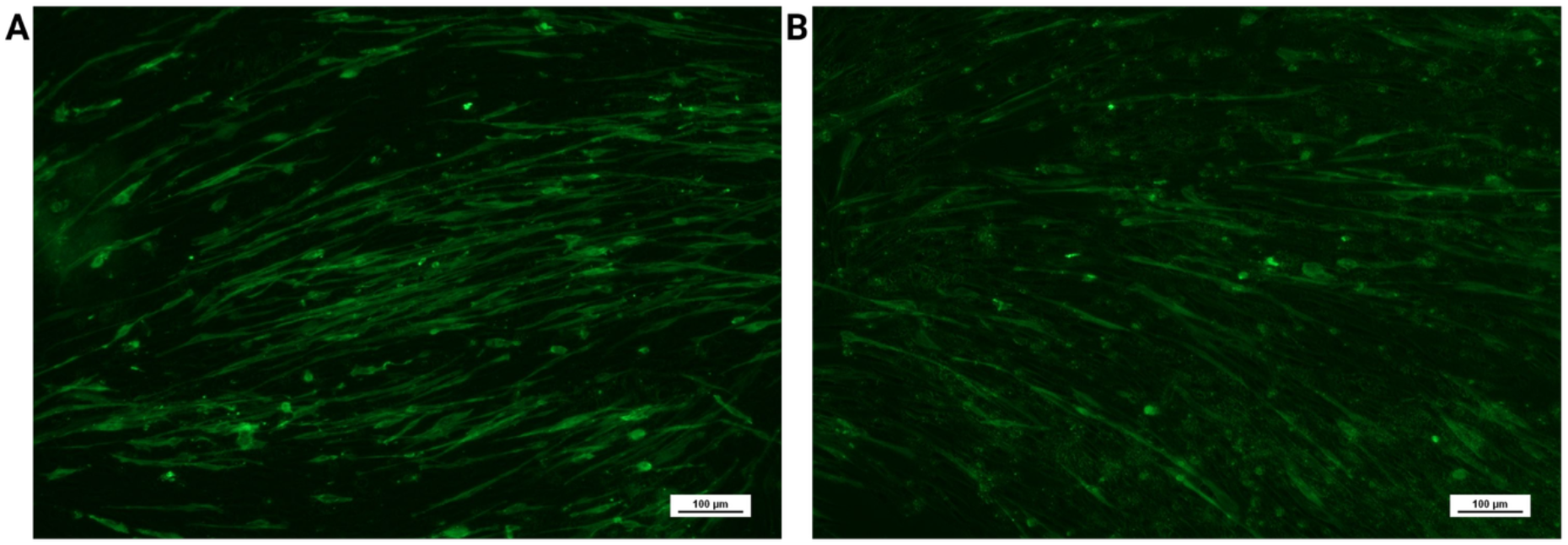
Immunocytochemical detection of sarcomeric α-actinin (SAA) in SsEC-derived myotubes on day 17 of differentiation. Representative fluorescence micrographs of SsECs differentiated under Condition 1 (C1; MN1/MN2) (A) or Condition 2 (C2; MS1/MS2) (B) show elongated, multinucleated myotubes with positive SAA staining. Scale bars = 100 µm.

**Figure 10.**
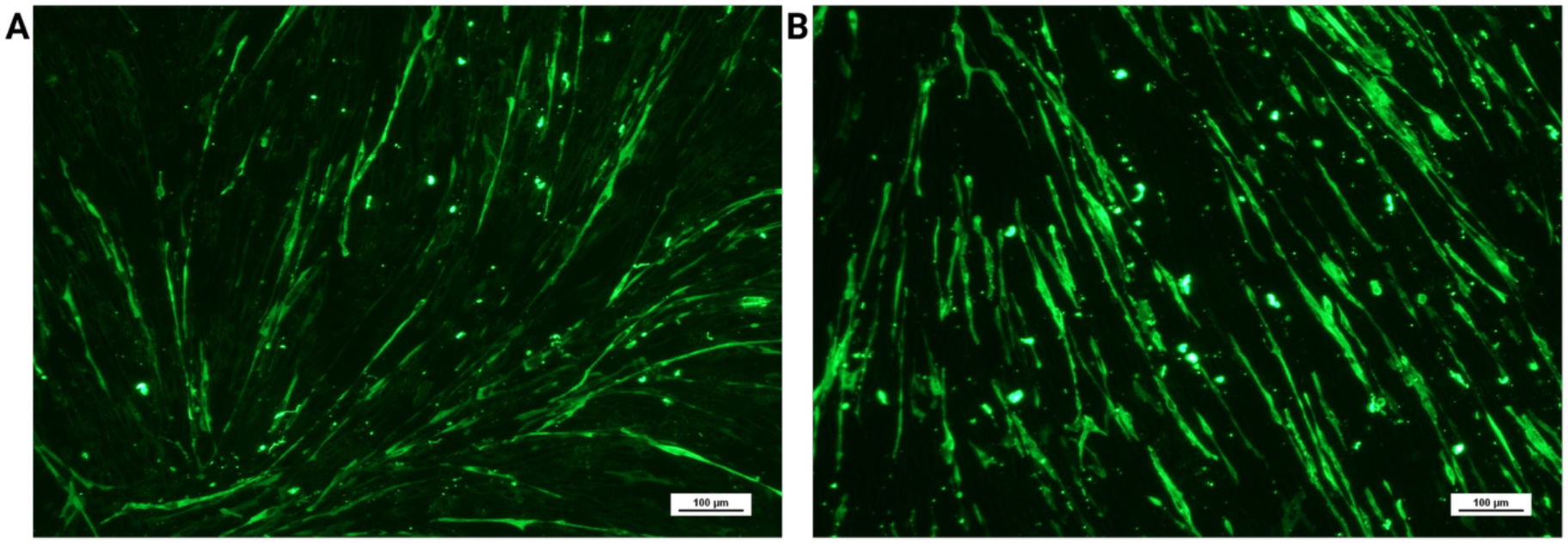
Immunocytochemical detection of myosin heavy chain (MHC) in SsEC-derived myotubes on day 17 of differentiation. Representative fluorescence micrographs of SsECs differentiated under Condition 1 (C1; MN1/MN2) (A) or Condition 2 (C2; MS1/MS2) (B) show elongated, MHC-positive myotubes. Scale bars = 100 µm.

### Blastula-derived ESC-like colonies require distinct ECM for attachment and expansion

Cells isolated from early (∼160 hpf) and mid-blastula (∼193 hpf) stage Atlantic salmon embryos displayed morphology and growth behavior that were clearly distinct from those of late embryonic SsEC cultures. When plated on laminin-coated surfaces, blastula-derived cells formed compact, tightly packed colonies with a high nucleus-to-cytoplasm ratio and smooth colony boundaries, consistent with an embryonic stem cell (ESC)-like morphology (Figure 11 A-F). Colonies became visible by approximately day 10 post-plating and were amenable to manual picking and expansion by approximately day 14.

**Figure 11.**
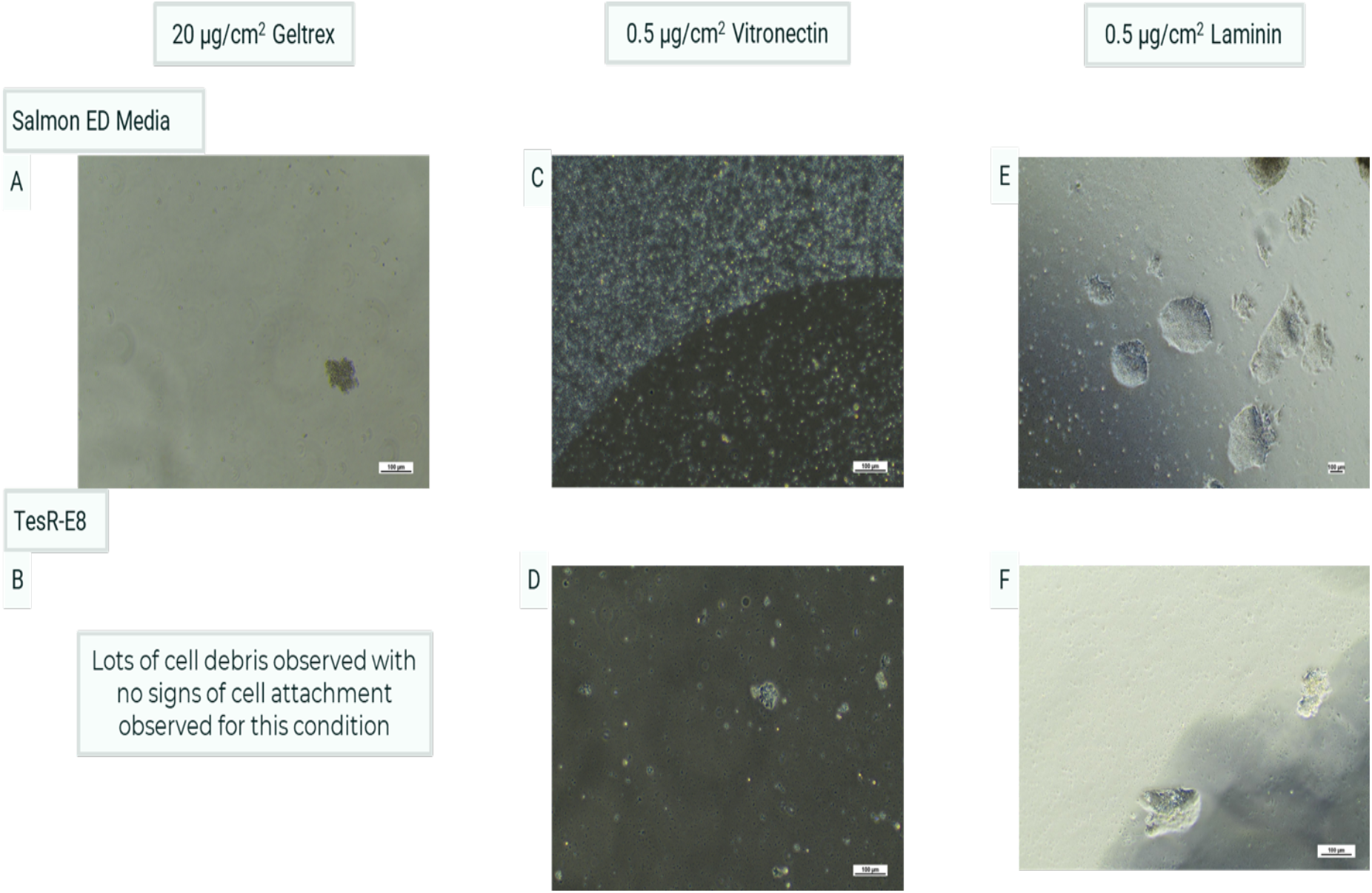
Derivation of cells from mid-blastula-stage Atlantic salmon (*Salmo salar*) embryos. Cells were assessed for attachment and growth on different extracellular matrix substrates under two culture conditions. No cell attachment or colony formation was observed on Geltrex or vitronectin in either medium (A–D). In contrast, laminin supported attachment and the formation of compact, colony-forming populations with embryonic stem cell-like morphology under both culture conditions (E, F).

In contrast to their robust growth on laminin, blastula-derived cells failed to attach or form colonies when cultured on vitronectin or Geltrex under otherwise identical conditions. In these contexts, cells remained largely non-adherent or exhibited only transient attachment without sustained proliferation or colony formation. This strict substrate dependence contrasted sharply with the behavior of SsECs derived from later embryonic stages, which preferentially attached to and expanded on vitronectin but did not form compact colonies characteristic of blastula-stage cultures.

Together, these observations demonstrate that embryonic stage of origin strongly influences both cell morphology and extracellular matrix requirements in Atlantic salmon embryonic cultures. Whereas blastula-derived cells exhibit an ESC-like colony phenotype that is selectively supported by laminin, late embryonic progenitors (SsECs) display a more elongated, progenitor-like morphology and require vitronectin for long-term attachment and expansion.

## Discussion

In this study, we report the establishment and comprehensive characterization of an embryonic-derived Atlantic salmon muscle progenitor cell line (SsEC) that exhibits robust long-term expansion and efficient, controllable myogenic differentiation under defined *in vitro* conditions. Stable muscle progenitor cell lines from teleost species remain limited, with many existing systems relying on short-lived primary cultures or heterogeneous populations with restricted differentiation fidelity and extensive donor-to-donor variability (Gabillard et al. 2010; Saad et al. 2023). In contrast, SsECs could be serially expanded for more than 30 passages while maintaining molecular and functional features consistent with a lineage-restricted myogenic progenitor, addressing a key technical bottleneck in fish muscle cell biology and aquaculture-relevant *in vitro* models.

A defining feature of SsEC derivation was a stringent and reproducible requirement for vitronectin as an extracellular matrix (ECM) substrate. While these embryonic-derived cells exhibited transient attachment to gelatin or laminin, only vitronectin supported sustained proliferation and long-term passaging. ECM-dependent regulation of progenitor behavior is well described in mammalian systems, where integrin-mediated interactions with defined matrices critically influence self-renewal and lineage commitment. However, ECM requirements in teleost embryonic progenitors remain comparatively underexplored. Our data demonstrate that ECM dependence in Atlantic salmon embryonic cells is both stage-specific and functionally consequential, with vitronectin selectively supporting late embryonic mesoderm-derived progenitors, whereas laminin was permissive only for earlier blastula-derived cultures (Johnston et al. 2011; Musialak et al. 2023).

Bulk RNA-sequencing provided strong molecular evidence that SsECs represent a stable, lineage-restricted myogenic progenitor population rather than a heterogeneous embryonic cell mixture. Principal component analysis demonstrated clear segregation of SsECs from the non-myogenic Atlantic salmon kidney (ASK) cell line, with minimal transcriptomic divergence between early and late passage SsECs. This stability is notable, as transcriptional drift during extended *in vitro* culture is a common limitation of fish cell lines. Differential expression analysis revealed significant enrichment of genes associated with myogenesis, sarcomere organization, and cytoskeletal architecture in SsECs, whereas ASK cells preferentially expressed endothelial- and kidney-associated transcripts. These transcriptomic profiles confirm that SsECs maintain a coherent muscle progenitor identity during prolonged expansion.

Targeted RT–qPCR analyses corroborated these findings and further demonstrated that lineage fidelity is preserved across passages. Sustained or elevated expression of the myogenic progenitor marker myf5 at later passages indicates maintenance of progenitor identity, while reduced expression of terminal muscle markers (acta1, tmp1, myl1) at higher passage suggests limited spontaneous differentiation during expansion. Such properties are highly advantageous for controlled differentiation protocols and distinguish SsECs from many previously reported teleost muscle cultures, which frequently exhibit premature differentiation or lineage instability (Gabillard et al. 2010; Saad et al. 2023). These observations align with conserved roles of myogenic regulatory factors and satellite cell-associated transcriptional programs in vertebrate muscle development and regeneration (Sultan et al. 2021; Ramírez de Acuña et al. 2022).

Functionally, SsECs demonstrated robust myogenic competence when subjected to a two-step differentiation protocol involving presomitic mesoderm-like induction followed by stage-specific modulation of growth factor signaling. This strategy mirrors differentiation frameworks developed for mammalian pluripotent and progenitor cells and leverages conserved developmental signaling logic, including WNT activation and TGF-β/BMP inhibition (Sanaki-Matsumiya et al. 2024). Under these conditions, SsECs formed multinucleated, sarcomere-bearing myotubes expressing myosin heavy chain and sarcomeric α-actinin, demonstrating that transcriptional activation of myogenic programs was accompanied by structural maturation and assembly of contractile machinery. This functional outcome establishes SsECs as a practical system for both mechanistic studies of teleost myogenesis and applied investigations relevant to cellular aquaculture (Johnston et al. 2011; Joyce 2023; García-Pérez et al. 2025).

In parallel with SsEC derivation, we observed that cells isolated from earlier blastula-stage embryos displayed markedly different morphology, growth behavior, and ECM dependence. These blastula-derived cells formed compact, tightly packed colonies with high nuclear-to-cytoplasmic ratios and smooth colony boundaries when cultured on laminin, closely resembling embryonic stem-like (ESC-like) cell populations described in other teleost species. Fish ESC-like cells have been reported from blastula-stage embryos of medaka, zebrafish, and several marine and freshwater species, and are characterized by colony-based growth, prolonged self-renewal, and expression of pluripotency-associated markers, although their developmental potential and stability vary between species and derivation protocols (Hong and Schartl, 2006; Alvarez et al. 2007; Hong et al. 2011; Son and Gong 2022). Consistent with this literature, blastula-derived salmon cells in the present study failed to attach or expand on vitronectin or Geltrex, underscoring a distinct, stage-specific ECM requirement that contrasts with the behavior of later embryonic progenitors.

The divergent properties of blastula-derived ESC-like cultures and late embryonic SsECs highlight a developmental transition from early pluripotent or multilineage blastomeres to lineage-restricted mesodermal progenitors during Atlantic salmon embryogenesis. Similar stage-dependent shifts have been reported in other teleost systems, where blastula-stage cells display broad developmental competence and colony-based growth, whereas later progenitors adopt elongated morphologies, reduced plasticity, and ECM preferences aligned with their emerging tissue identities (Hong et al. 2011; Son and Gong, 2022). These findings emphasize that embryonic stage of isolation is a critical determinant of *in vitro* phenotype and must be carefully considered when developing fish cell models for both basic and applied research.

Future work will build on the experimental foundation established here by extending functional and molecular characterization of SsECs across additional biological contexts. In vivo transplantation assays will provide an opportunity to assess myogenic contribution and engraftment potential within host tissue, offering further insight into lineage potential and functional maturation. In parallel, deeper multi-omics approaches, including single-cell transcriptomics and epigenomic profiling, will help resolve regulatory networks governing progenitor maintenance, lineage commitment, and terminal differentiation. Together with the robust derivation strategy, transcriptomic stability, and reproducible myogenic differentiation demonstrated in this study, these avenues position SsECs as a versatile platform for advancing comparative developmental biology, applied aquaculture research, and emerging cellular seafood technologies (Macqueen et al. 2008; Trace et al. 2024).

## Supplementary Data

**Table S1:**
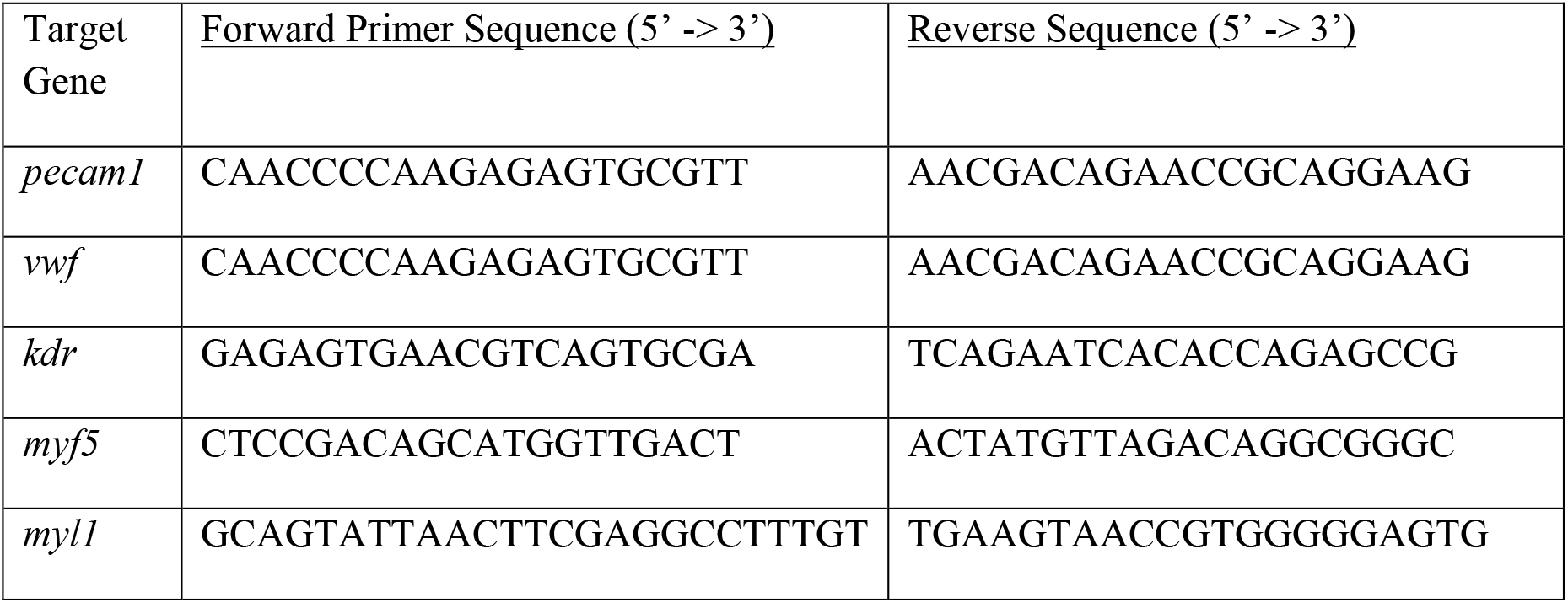
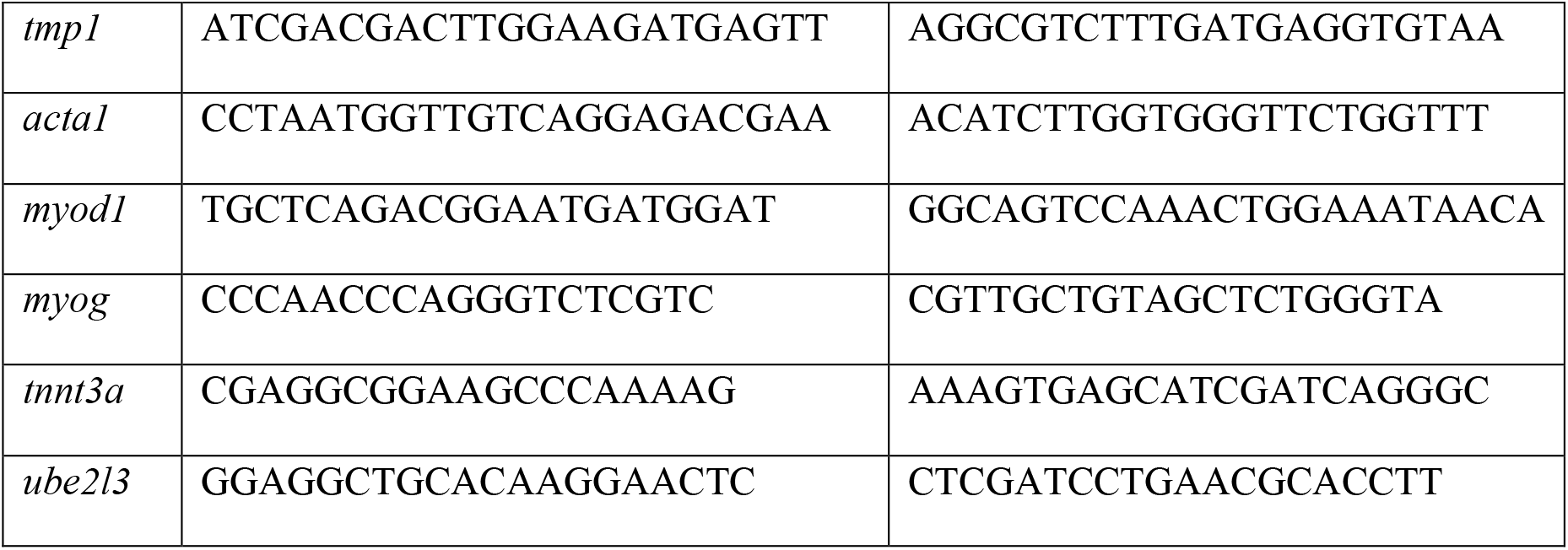
Primers for detecting *ube2l3, myf5, myod1, myog, tnnt3a, myl1, tmp1, acta1, vwf, pecam1*, and *kdr* gene expression using qPCR.

## Data Availability Statement

The datasets used and analyzed during the current study are available to download from Roslin Technologies servers on reasonable request from the corresponding author.

## Ethics Statement

Fertilized Atlantic salmon ova were obtained from a licensed supplier. Embryos were handled in accordance with institutional and national guidelines for the use of fish embryos in research. All work was conducted in accordance with institutional and national guidelines governing the use of fish embryos and did not require regulated animal procedures.

## Author Contributions

KN and SW performed cell derivations, culture optimization and differentiation. PJM/KN conceived the study which was supervised by PJM. KN and PJM drafted the manuscript with input from all authors. DR oversaw the project and provided scientific input. All authors approved the final manuscript.

## Acknowledgments

We thank colleagues at Roslin Technologies staff for discussions and technical assistance. We are grateful to Benchmark Genetics especially Professor Ross Houston for provision of fertilized ova and helpful discussions.

